# Soil properties in agricultural systems affect microbial genomic traits

**DOI:** 10.1101/2024.10.23.619851

**Authors:** Tim Goodall, Susheel Bhanu Busi, Robert I. Griffiths, Briony Jones, Richard F. Pywell, Andrew Richards, Marek Nowakowski, Daniel S. Read

## Abstract

Understanding the relationships between bacterial taxa, their ecological and genomic traits, and their environment, is important for elucidating the mechanisms that drive microbial community dynamics and their roles in ecosystem functioning. This is especially true for soils, where dramatic shifts in resource input or physicochemical properties occur through land use and agricultural practices. Here, we examined the relationships between soil properties and bacterial traits within highly managed agricultural soil systems subjected to arable crop rotations or management as permanent pasture. We assessed the bacterial communities within these soils using amplicon sequencing and assigned each amplicon trait scores for rRNA copy number, genome size, and GC content, which are classically associated with potential growth rates and specialisation. We also calculated the niche breadth trait of each amplicon as a measure of social ubiquity within the examined samples. Within this soil system, we demonstrated that pH was the primary driver of bacterial traits. The weighted mean trait scores of the samples revealed that bacterial communities associated with soils at lower pH (<7) tended to have larger genomes (possess more potential plasticity), have more rRNA (higher growth rate potential), and are more ubiquitous (have less niche specialisation) than the bacterial communities from higher pH soils. Our findings highlight not only the association between pH and bacterial community composition but also the importance of pH in driving community functionality by directly influencing genomic and niche traits.

## Introduction

Microbial genomic traits such as genome size and rRNA copy number are key indicators of niche breadth and ecological strategies in soil bacterial communities^1^. These traits reflect the ability of microorganisms to adapt to varying environmental conditions^2^, including nutrient availability, pH, and organic matter content. Understanding the relationship between soil properties and microbial genomic traits is crucial for advancing our knowledge of microbial ecology, particularly in agricultural systems, where soil management practices can significantly alter these properties.

Agricultural soils represent a significant portion of land use in the United Kingdom, with 70% of the total land area classified as Utilised Agricultural Area (UAA)^3^. In 2023, the total arable area in the UK was reported to be just over 6.0 million hectares, accounting for approximately 36% of the UAA^3^. Soils in these agricultural systems vary widely in pH and organic matter content, which are critical factors that influence soil dwelling microbial communities and their genomic traits^4,5^. Previous studies have shown that conversion to arable land leads to modified microbial communities, where the consequences include reductions in functionality and decreases in genes relating to important biogeochemical cycles, including those that indlfluence the cycling and fate of carbon, phosphorous and nitrogen containing compounds^6^. Simultaneously, the age of managed and permanent pastures, key for providing food and income, affects the microbial community, especially bacteria, through the influence of soil physicochemical properties, such as pH^7^.

Microbial metabolic versatility^8^ and the ability of taxa to thrive in diverse and fluctuating environments^9^ are typically associated with larger genome sizes and increased rRNA copy numbers. Conversely, smaller genomes and fewer rRNA copies suggest specialisation and efficiency under more stable or resource-limited conditions^2^. The ability of soil microbial communities to adapt to changes in soil properties, such as pH and organic matter content, is reflected in these genomic traits, which in turn influence the overall health and functionality of the soil ecosystem. For example, Wilhelm et al.^10^ demonstrated a positive relationship between genome size and rRNA copy number in soil bacteria. These genomic traits are relevant for understanding the classical concept of ecological niche breadth, which defines the range of conditions under which organisms can survive in^11^. Although niche breadth has previously been assessed with respect to environmental variables, it has been limited to specific taxa^12^ or environments^13^. Importantly, the notion of generalists, taxa found in many samples or predefined habitats, and specialists, taxa that are rare or highly fastidious^14^, have not been assessed in soils under land use effects, necessitating the need to understand how anthropogenic influences affect the social niche breadth of microorganisms.

To test the relationship between microbial traits and environmental influences, we assessed microbial communities and their traits in soils from arable and permanent grassland across 245 samples from 68 fields at 20 farms in southern UK/England. By leveraging systematic sampling and molecular methods, our study provides novel insights into the relationship between soil properties and microbial genomic traits in agricultural systems, especially arable and permanent pastures, the latter of which are relatively unknown. Our findings highlight the importance of pH as a key factor influencing microbial genomic traits, such as size and niche breadth, challenging the view that nutrient availability (carbon content) is a key determinant of these traits. These results have important implications for understanding microbial adaptation in agricultural soils and the management of soils in these systems for specific ecological outcomes.

## Methods

### Sample collection

During the Autumn of 2019, 245 subsurface (<15 cm) soils were sampled from 68 fields in southern England, by Agrii-Masstock Arable (UK) Limited. The samples were collected into sample boxes and transported immediately to NRM – Cawood Scientific (Berkshire, UK) for processing.

### Physicochemical measurements and DNA extraction

Soil physicochemical measurements were made at NRM – Cawood Scientific (Berkshire, UK): pH, organic matter content as a percentage of dry mass by loss on ignition (LOI); phosphorus, potassium, magnesium as mg per L; and sand (0.063 to 2.000 mm), silt (0.002 to 0.063 mm), and clay (<0.002 mm) content as a percentage of total. The sample properties are listed in Supplementary Table 1.

DNA extraction was performed on 0.2 g of dried, milled soil sample using the Qiagen Power Soil HTP-96 kit (Cat. No. 12955-4) following manufacturer’s instructions.

### PCR and sequencing

The extracted DNA was used to generate metabarcoding sequence data. Briefly, amplicons were produced using a 2-step amplification approach, with Illumina Nextera tagged primers targeting the V4-5 region of 16S rRNA: 515f GTGYCAGCMGCCGCGGTAA and 806r GGACTACNVGGGTWTCTAAT^15^ with each primer modified at 5’ with the addition of Illumina pre-adapter and Nextera sequencing primer sequences.

Amplicons were generated using high-fidelity DNA polymerase (Q5 Taq; New England Biolabs). After an initial denaturation at 95 °C for 2 min, the PCR conditions were as follows: 30 cycles of denaturation at 95 °C for 15 s, annealing at 55 °C for 30 s, and extension at 72 °C for 30 s. A final extension step of 10 min at 72 °C was performed. PCR products were purified using Merck MultiScreen PCR filter plates following manufacturer’s instructions. MiSeq adapters and 8nt dual-indexing barcode sequences^16^ were added during the second PCR amplification step. After an initial denaturation at 95 °C for 2 min, the PCR conditions were eight cycles of denaturation at 95 °C for 15 s, annealing at 55 °C for 30 s, and extension at 72 °C for 30 s, with a final extension of 10 min at 72 °C.

The amplicon size was determined using an Agilent 2200 TapeStation system, and the library was normalised using the NGS Normalization Kit (Norgen Biotek), with subsequent quantification using the Qubit dsDNA HS kit (Thermo Fisher Scientific). The pooled libraries were further purified by gel extraction (QIAquick, Qiagen) and diluted to 400pM with 7.5% Illumina PhiX. Denaturation of the library was achieved by adding a 10% final volume of 2N NaOH and incubating at room temperature for 5 min, followed by neutralisation with an equal volume of 2N HCl. The library was then diluted to its load concentration using Illumina HT1 Buffer. The final denaturation was performed by heating to 96°C for 2 min, followed by cooling in crushed ice. The amplicon library was sequenced on an Illumina MiSeq using V3 600 cycle reagents.

### Bioinformatic analysis

Illumina demultiplexed sequences were processed in R using Dada2^17^ to filter, denoise, and merge the sequences with the following parameters: primer sequences were removed with cutadapt^18^ and reads were truncated to 250 and 200 bases, forward and reverse, respectively. The filtering settings were as follows: maximum number of Ns (maxN) = 0 and maximum number of expected errors (maxEE) = (5,5). Sequences were dereplicated, and the DADA2 core sequence variant inference algorithm was applied. Forward and reverse reads were then merged using the *mergePairs* function to produce amplicon sequence variants (ASVs). Chimeric sequences were removed using *removeBimeraDenovo* at default settings and sequence tables were constructed from the resultant ASVs. ASVs were subjected to taxonomic assignment using the *assignTaxonomy* function and SILVA v138.1^19^ training database.

### Bacterial community beta-diversity and environmental drivers

Examination of community composition was carried out in R using Microeco^20^ and Vegan^21^. Briefly, samples with less than 8,000 reads were removed along with the removal of sequence variants identified as chloroplast and mitochondrial, or those not belonging to the domains of Bacteria or Archaea, and each sample’s data were rarefied to 7,555 reads before Bray–Curtis distance matrix calculations and non-metric multidimensional scaling (NMDS) analysis and plotting. Correlations between environmental variables were examined at the phylum level using the *cal_cor* function.

### Genomic trait assessment

To estimate the genome size of the identified taxa based on taxonomic information, ASVs were assigned values using representative genomes present in the IMG-ER public database^22^. To do this ASV taxonomies were matched iteratively to IMG-ER representative genomes as described by Markowitz et al.^22^. Genomic traits summarised by the database were downloaded (September 20^th^, 2024) for all isolate, single-cell amplified, and metagenome-assembled genomes. Based on the availability of taxonomies in the IMG-ER database, relevant genomic traits such as genome size, rRNA operon copy number, coding density, and GC content were retained. ASVs with unclassified genera were iteratively matched to higher taxonomic ranks. For example, if a particular genus did not have a genome size, the genome size of the family to which the genus was ascribed was used. To confirm rRNA copy numbers, a FASTA file containing the representative sequences of all ASVs was used to approximate rRNA copies using an Artificial Neural Network Approximator (ANNA16)^23^. A consensus of the rRNA copies was used for subsequent analyses. The weighted mean rRNA copies, weighted mean GC, and weighted mean genome size of each sample were calculated using the rarified sequence abundance table.

### Social niche breadth

To determine the niche breadth of the identified ASVs within the dataset, we used the social niche breadth (SNB) metric, as described by Miejenfeldt et al.^1^. SNB quantifies the diversity of ecological interactions that a microbial taxon engages in by analysing co-occurrence patterns across samples. First, the taxonomic profiles were rarefied to equal sequencing depths to avoid bias from differential sequencing coverage. The rarefied data were then formatted and processed using the *calculate_SNB.py* script from the SNB analysis toolkit (https://github.com/MGXlab/social_niche_breadth_SNB). SNB scores for each ASV were calculated based on their co-occurrence with other taxa, reflecting the range of ecological networks an ASV interacts with, where higher scores indicate a more generalist strategy and lower scores reflect specialisation. For each sample, a weighted mean SNB score was computed by combining the SNB scores of all ASVs present, weighted individually by their abundance (count), to provide an overall measure of the sample’s community niche breadth. This approach was used to assess how environmental factors influence microbial generalisation or specialisation across samples.

### Phylogenetic and PhyloFactor analysis

For phylogenetic analysis, we generated a multiFasta file from the amplicon sequence variant (ASV) table produced by the DADA2 pipeline. We utilised the DECIPHER package in R to align the DNA sequences. Following alignment, we constructed a Neighbor-Joining (NJ) tree based on a distance matrix calculated from the aligned sequences using the maximum likelihood (ML) method, employing the General Time Reversible (GTR) model for optimisation. The resulting optimised ML tree served as the basis for subsequent phylogenetic analyses, including downstream PhyloFactor analysis^24^. This analysis incorporated taxonomic data, which were combined into a single column (lineage), along with the ASV abundances and associated metadata. To further evaluate potential biases in the estimated SNB across different lineages, including dominant and rare taxa, we assessed phylogenetic signals within the dataset. Specifically, we estimated Blomberg’s K and Pagel’s Lambda using phylogenetic generalised least squares (PGLS) regression for statistical inference (see Code availability).

As highlighted above, to analyse phylogenetic factors driving patterns in community composition, we employed the PhyloFactor package in R, where the input was the phyloseq object. We ensured that the phylogenetic tree was unrooted using the unroot function from the ape package, a critical step for the proper functioning of the PhyloFactor method. The ASV abundances were modelled as a response variable to the SNB score of the individual ASVs, applying a log-ratio transformation and establishing a significance threshold of 0.01. We specified several factors (n=15) to be evaluated, from which the detailed summaries of the PhyloFactor analysis were extracted, selecting for significant taxa.

For data visualisation, isometric log-ratio (ILR) boxplots, incorporating the ILR results into the phyloseq object through a custom function (see Code availability), were generated. This allowed for ILR stratification by high- or low-SNB scores, with separation based on median average weighted SNB scores. A phylogenetic tree was constructed to visualise the relationships between taxa and the SNB category, highlighting the significant taxa identified in the analysis (see Code availability).

### Data analysis and figures

R (v4.1) was used for data analysis and figure generation using the *ggplot2* package. The *mgcv* package was used for the regression analyses, while any differential analysis was performed using functions in base R.

## Results

### pH and LOI together shape the microbial community composition

Arable and permanent pastures have been shown to support the functioning of varied microbial communities. To understand how arable and permanent pastures vary in microbial composition, diversity, and abundance, in this study, we analysed 245 soil samples collected from arable and permanent pasture fields across southern England (Figure 1a).

**Figure 1.**
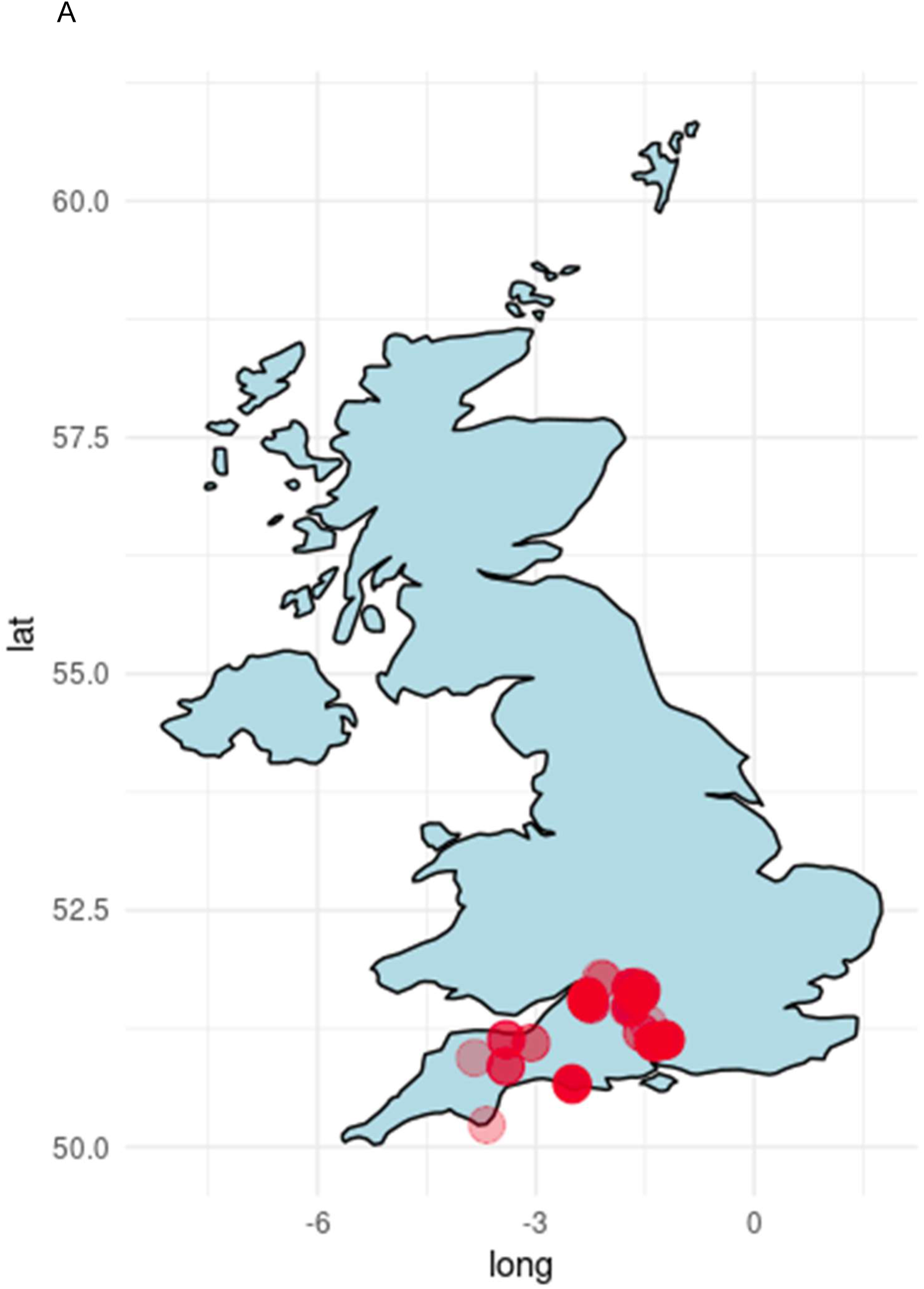
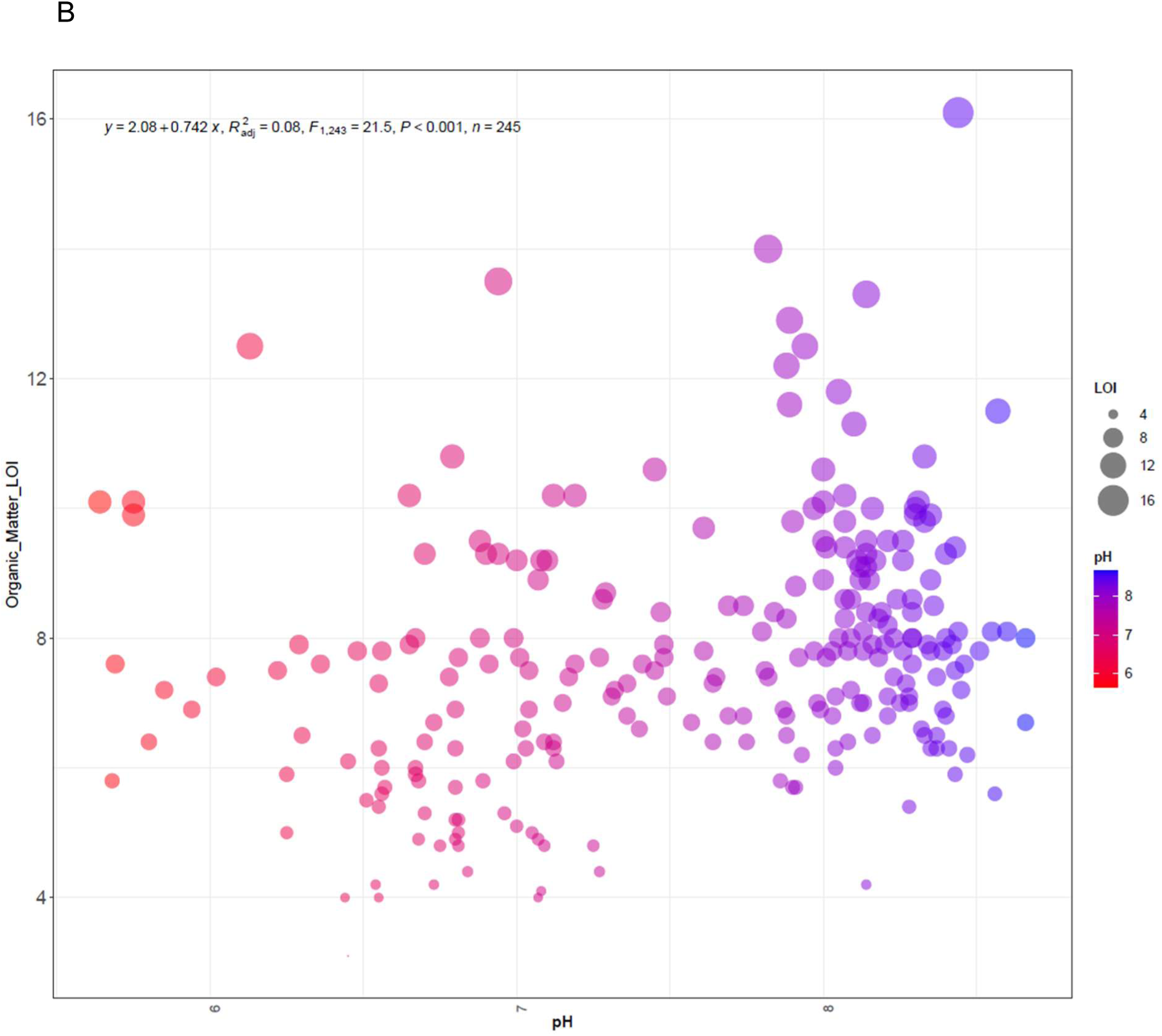
**(a)** Map of the United Kingdom showing the sample site distribution within southern England (n=245; point alpha = 0.3); **(b)** Relationship between soil samples’ organic matter content (LOI) and soil pH (*p* < 0.001, linear regression). The size of the filled circles indicates the measured LOI of individual samples.

The average pH of these soil samples was 7.6 (range 5.6 to 8.7), and the average LOI was 7.7% (range 3.1% to 16.1%). A weak but significantly positive relationship between pH and LOI (r² = 0.08, p < 0.001) was observed (Figure 1b), demonstrating that this sample set does not conform to the broadly typical trend of high LOI at low pH^25^. This non-conformity can be explained due to the limited land use range, and because of the prevalence of samples with basic parent material where this trend breaks down^26^. An ordination analysis (Figure 2) revealed that, as per previous studues^27^, the soil bacterial communities were primarily structured by pH.

**Figure 2.**
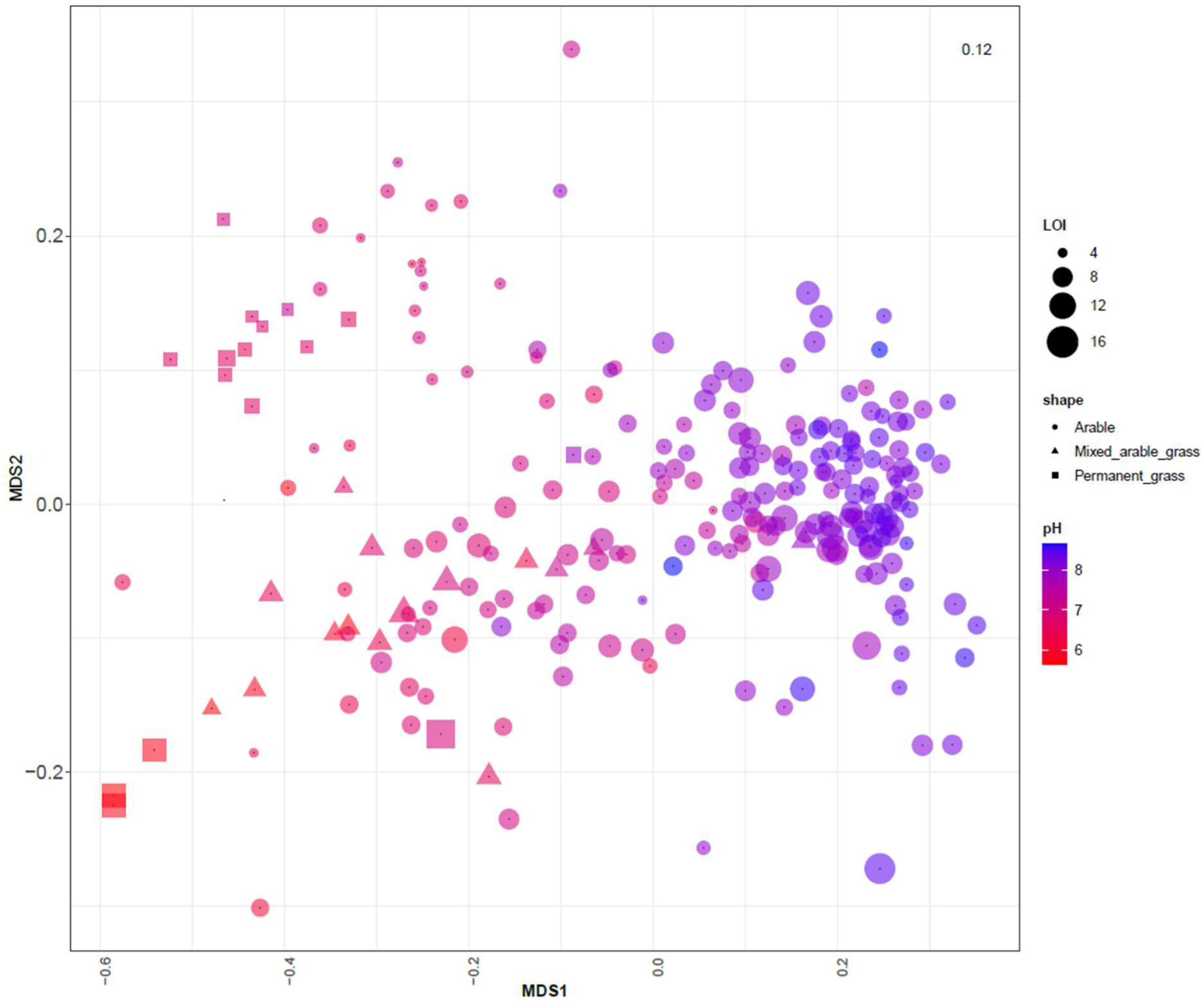
Non-metric multidimensional scaling (NMDS) plot of soil beta diversity, where point size indicates the percentage of organic matter by loss on ignition (LOI), point colour indicates soil pH, and shape indicates cropland rotation type. NMDS stress value (0.12) is indicated at the top right.

### Broad taxonomic variability across arable and permanent pastures

In-depth classification and analysis of the taxonomic components of the soils led to the observation of Actinobacteria, Proteobacteria, Chloroflexi, Verrucomicrobiota, Acidobacteria, Firmicutes, Bacteroidota, Crenarchaeota, Planctomycetota, and Myxococcota phyla being within the ten most abundant and cosmopolitan phyla within the soils (Figure 3a). Among these, the abundant phyla Entotheonellaeota, Crenarchaeota, Acidobacteria, Proteobacteria, Nitrospirota, NB1-j, Methylomirabilota, RCP2-54, and Bacteroidota were found to be positively correlated/associated with pH (Figure 3b). A selection of the same taxa also positively correlated with LOI. Conversely, taxa, such as Firmicutes, Verrucomicrobiota, and WPS-2, were negatively correlated with pH. Diversity (Simpson’s) was found to increase with soil pH (Figure 3b), again highlighting that these soils from highly managed land use exhibit atypical characteristics when compared to broader national surveys^27^.

**Figure 3.**
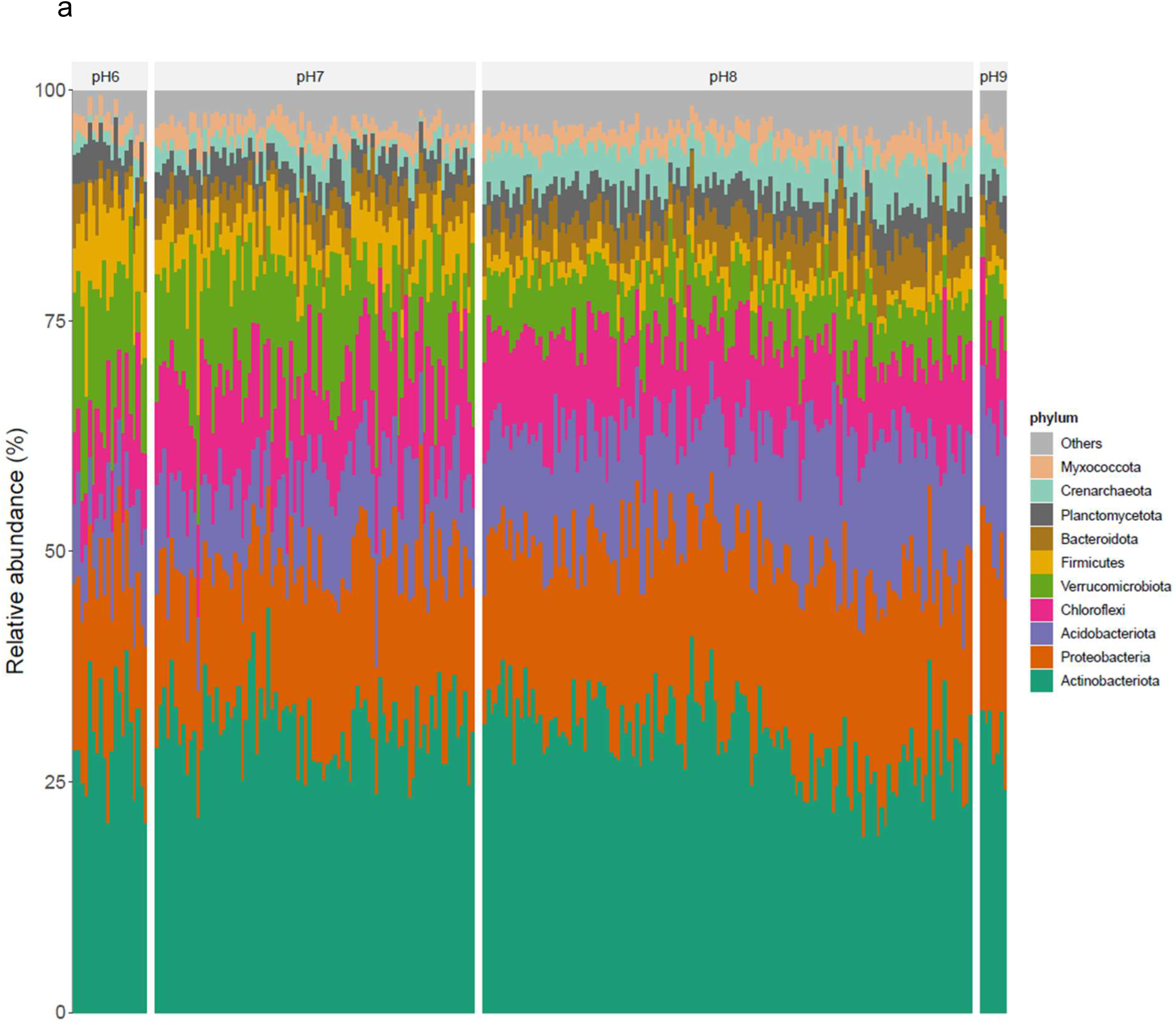
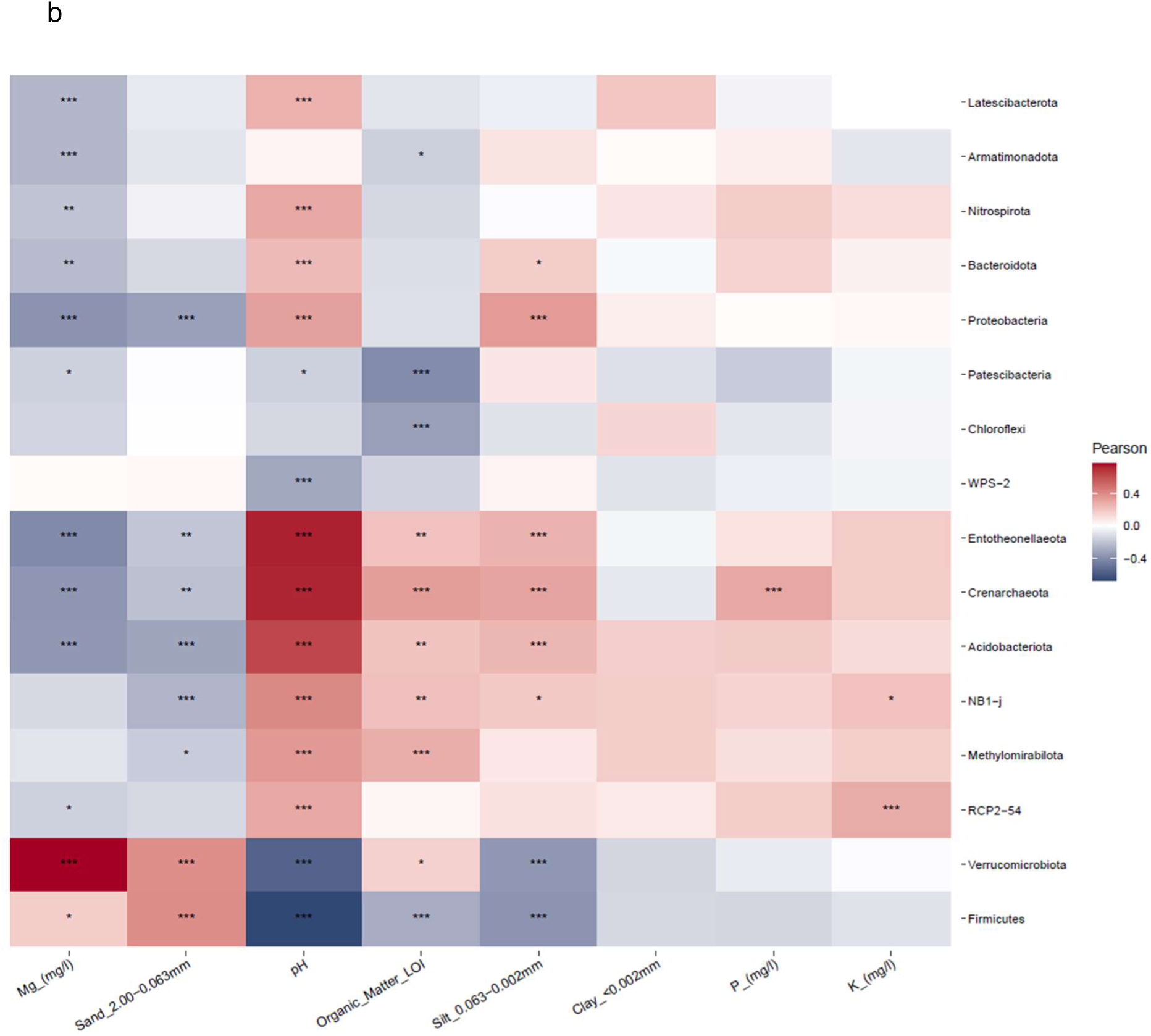
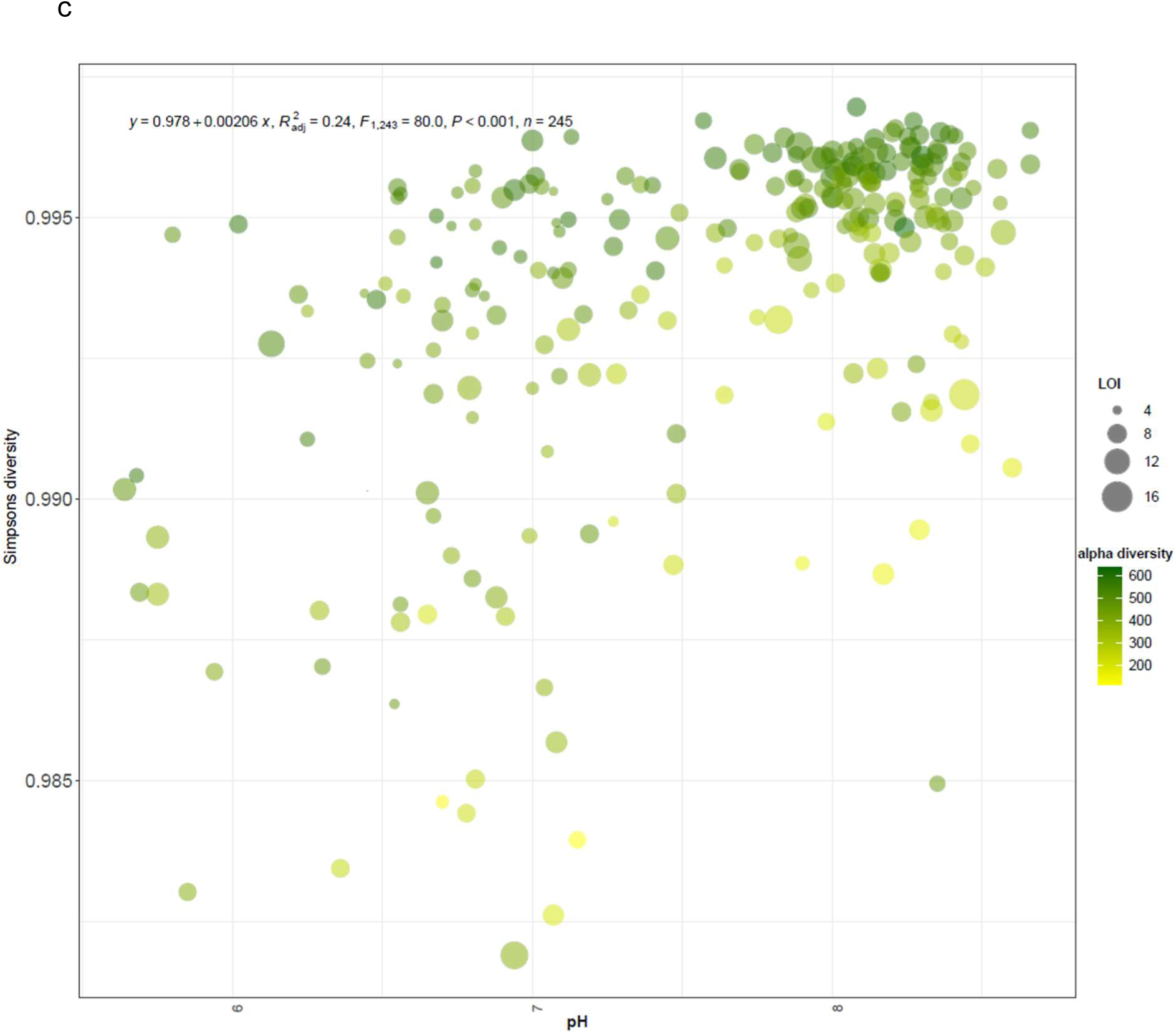
**(a)** bar plot of the top ten abundant phyla faceted by pH. pH6: samples ranging from pH 5.6-6.5, pH7: samples from pH 6.6-7.5, pH8: samples from pH7.6-8.5 and pH9: samples from and above pH 8.5. **(b)** correlations observed between all phyla and measured environmental variables (spearman’s correlation, p-values: ‘***’ 0.001, ‘**’ 0.01, ‘*’ 0.05). **(c)** Relationship between Simpson’s diversity index and soil pH (adj. R^2^ = 0.24, *p* <0.001, linear regression), LOI as the point size, and the sample’s alpha diversity as the colour gradient.

### Soil properties influence genomic traits of bacteria

Given the observed variability in community composition and abundance, it is important to understand the effect of inherent soil properties on bacterial communities within arable and permanent pastures. To achieve this, we assessed the relationship between the dominant soil properties (pH and LOI) and the genomic traits of bacteria found in these soils. Importantly, we focused on genome size and rRNA copy number which are indicative of life history, ecological strategies, and evolutionary adaptation. Genomic traits of the observed taxa were determined using genome representatives obtained from the publicly available IMG-ER database (*Methods*). We found a positive correlation between the weighted mean genome size and weighted mean rRNA copy number per genome (Figure 4). Interestingly, we found that several samples with larger mean genome sizes and increased rRNA copy numbers were observed at decreasing pH levels. Generalised additive model analysis (Table 1) confirmed mean average rRNA copy number (ave.rrna) and mean average genome size (ave.gs) are significantly associated with soil pH (adj. *p* < 0.05; Table 2).

**Figure 4.**
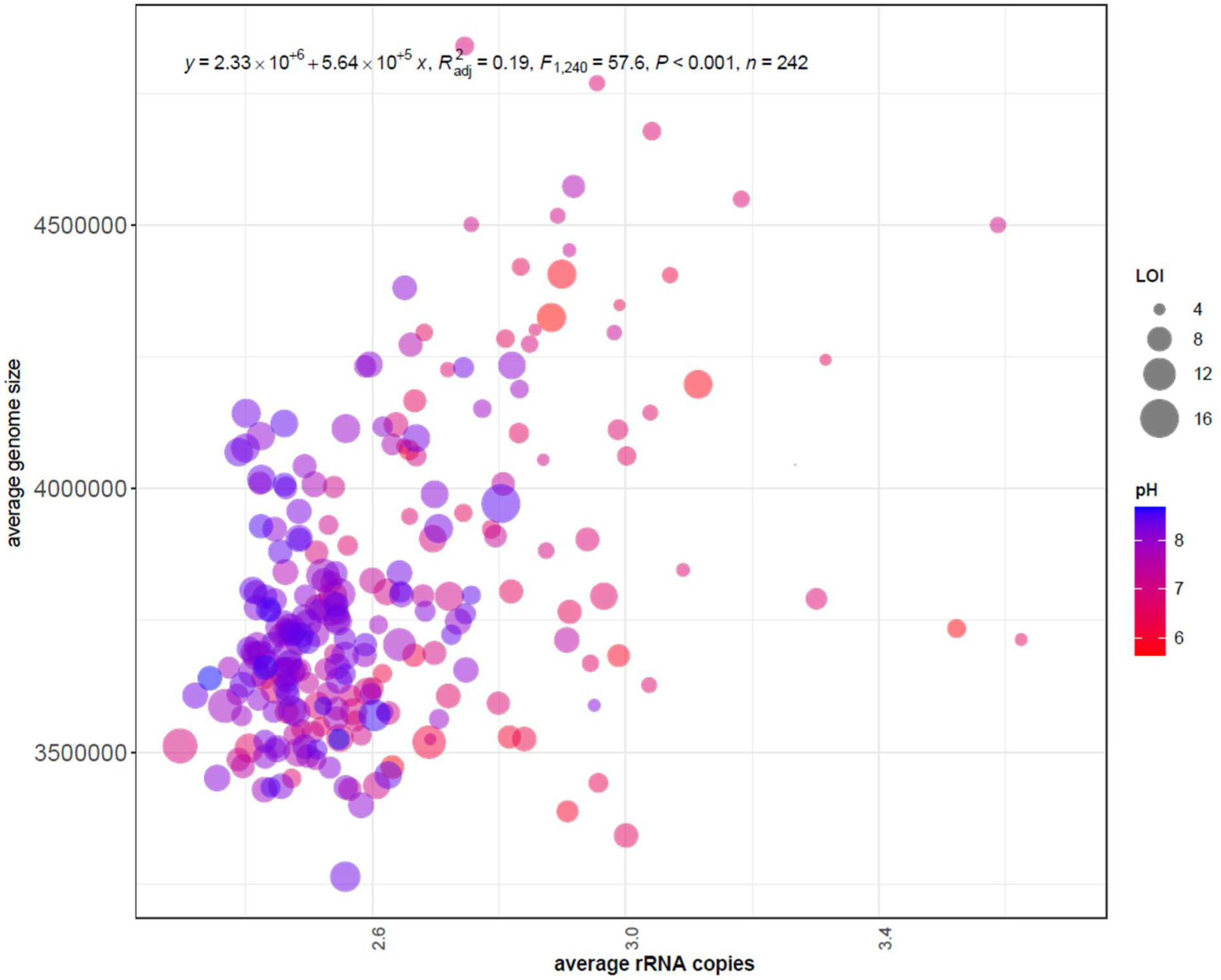
Scatter plot illustrating the significant positive relationship between the weighted mean genome size and weighted mean rRNA copy number, R^2^ = 0.19, *p* < 0.001, linear regression.; LOI as point size and sample pH as colour gradient

**Table 1.**
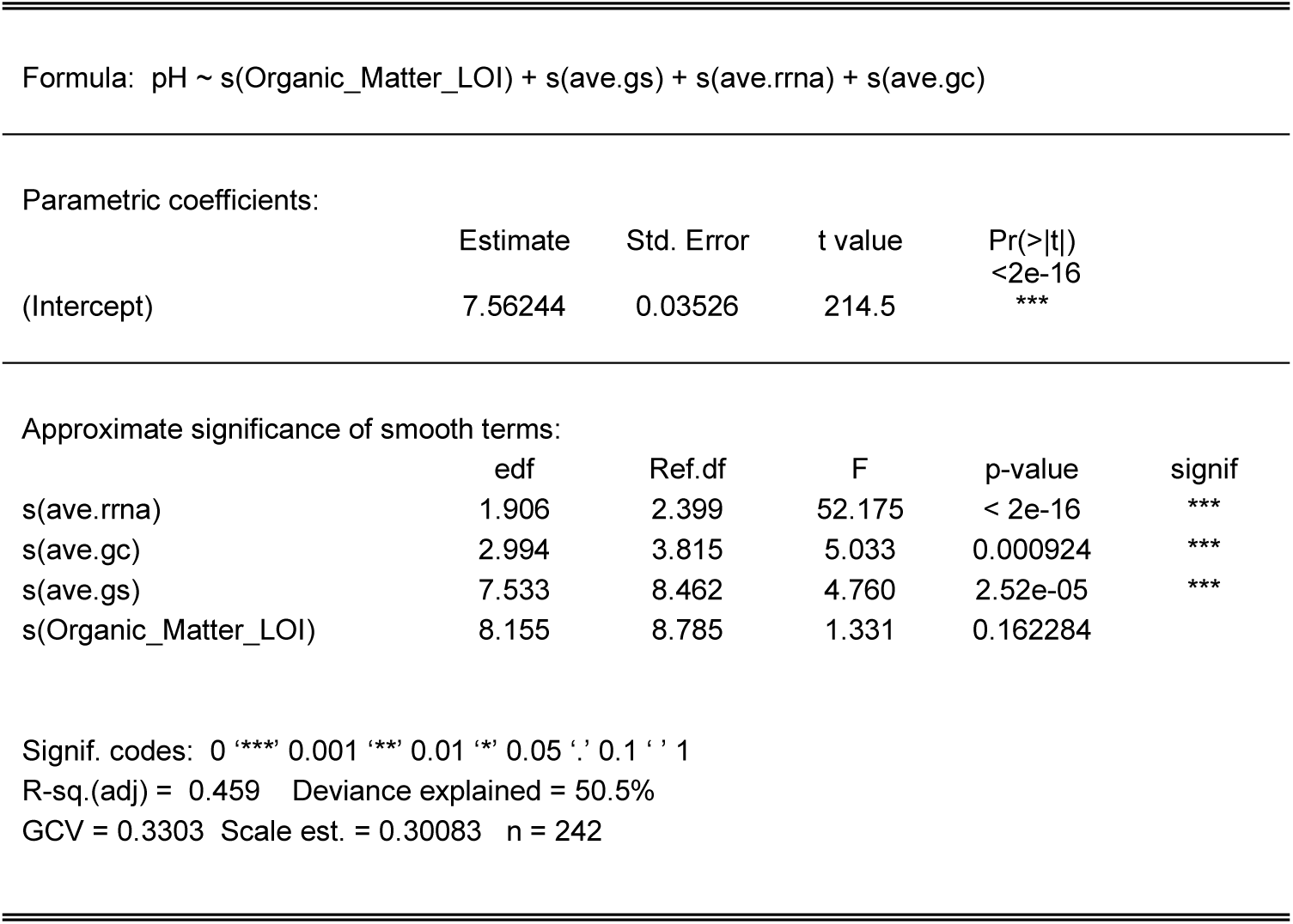
Summary of generalised additive model – significant relationships of weighted mean rRNA copies (ave.rrna), weighted mean GC content (ave.gc), and weighted mean genome size (ave.gs), with soil pH.

**Table 2.** Summary of generalised additive model – significant relationships of pH, LOI (Organic_Matter_LOI) and weighted mean rRNA copy number (ave.rrna), with weighted mean social niche breadth (ave.snb) score.

|  | Estimate | Std. Error | t value | Pr(>\|t\|) |
| --- | --- | --- | --- | --- |
| (Intercept) | 0.566325 | 0.0005157 | 1018 | <2e-16<br>*** |

Approximate significance of smooth terms:
|  | edf | Ref.df | F | p-value | signif |
| --- | --- | --- | --- | --- | --- |
| s(pH) | 7.869 | 8.663 | 74.332 | < 2e-16 | *** |
| s(Organic_Matter_LOI) | 1 | 1 | 11.912 | 0.000668 | *** |
| s(ave.rrna) | 6.771 | 7.866 | 5.98 | 1.53E-06 | *** |
| s(ave.gc) | 2.884 | 3.682 | 2.937 | 0.032546 | * |
| s(ave.gs) | 1 | 1 | 2.212 | 0.138335 |  |

Signif. codes: 0 '\*\*\*' 0.001 '\*\*' 0.01 '\*' 0.05 '.' 0.1 ' ' 1
R-sq.(adj) = 0.865 Deviance explained = 87.6%
GCV = 7.0331e-05 Scale est. = 6.4367e-05 n = 242

### Bacterial niche breadth is influenced by soil properties

Since microbial composition and genomic traits were influenced by soil properties, we used the social niche breadth (SNB) of the individual taxa to calculate each sample’s weighted mean SNB score. We found a clear distinction between the weighted mean SNB values, driven primarily by pH (Figure 5). For example, at low pH levels, samples exhibited a higher SNB score, whereas the score was reduced at higher pH. The genome sizes within these communities varied and were diverse, with average genome sizes ranging from 3,263,535 bp to 4,840,393 bp (3,805,988 bp average). To rule out phylogenetic and abundance biases, we applied phylofactorization to assess phylogenetic signals linked to SNB scores. Of the 15 factors tested, 46% (7/15) were associated with high SNB, whereas the remainder were associated with low SNB (Supplementary Fig. 2). Notably, these SNB scores were independent of phylogenetic classification (Supplementary Fig. 3), suggesting that environmental factors primarily shape community composition. Factor 6 was representative of all factors where taxa across the bacterial tree were included in the low and high SNB clades along the tree (Supp. Fig. 2). Additionally, phylogenetic and abundance factors did not bias SNB, as both dominant and rare taxa were represented in the low and high SNB groups (Supp. Fig. 2). To further investigate the drivers of SNB, that is, soil properties, we used generalised additive models to establish their importance and significance. Generalised additive model analysis confirmed that alongside pH and LOI, both average GC content (ave.gc; Supp. Fig. 1), and average rRNA copy number (ave.rrna) were significant drivers of SNB within the observed microbial community (adj. *p* < 0.05; Table 2).

**Figure 5.**
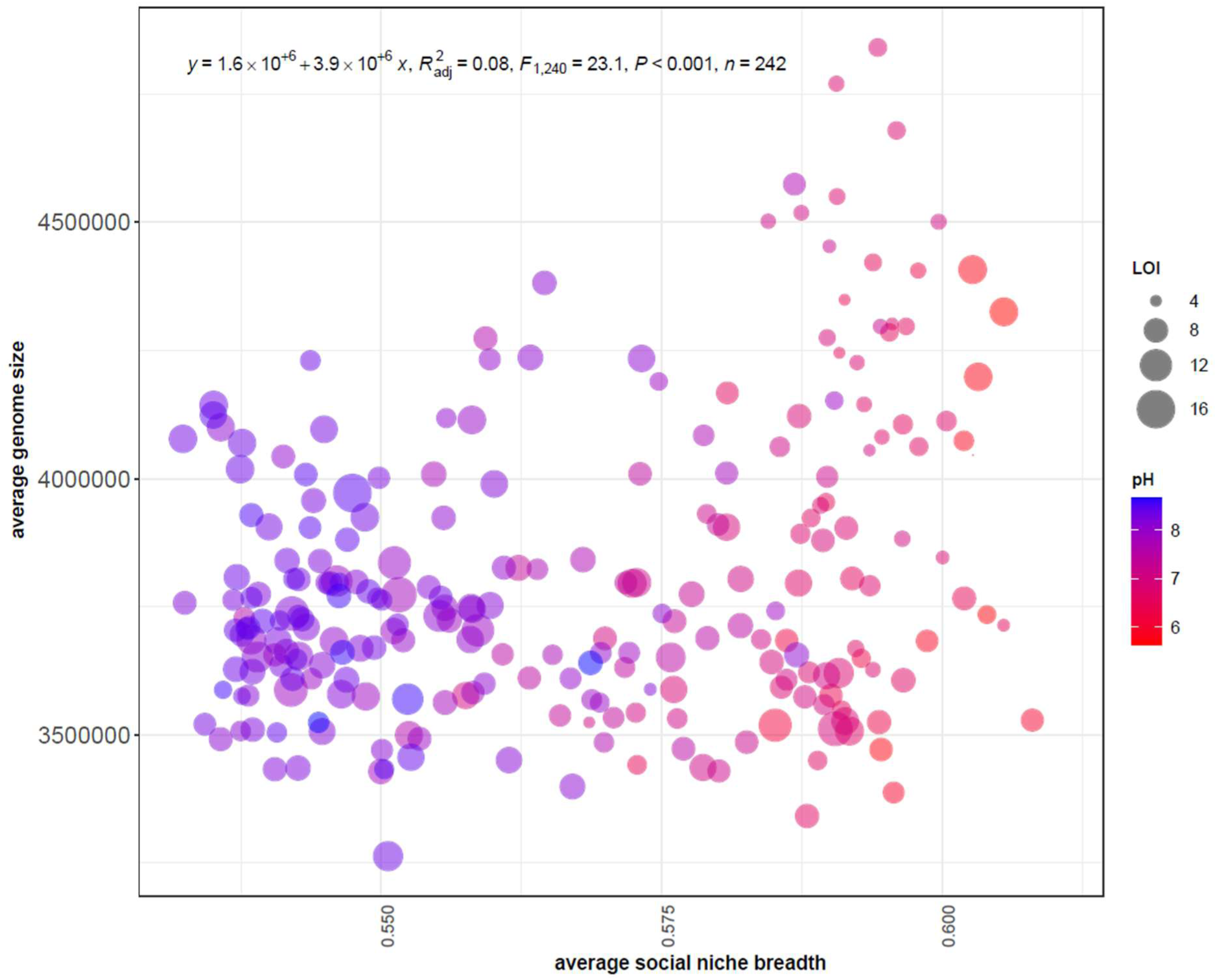
Scatter plot showing the positive relationship between the weighted mean genome size and weighted mean social niche breadth, R^2^ = 0.08, *p* < 0.001, linear regression. The pH colour gradients showed an increasing mean social niche breadth and mean genome size with acidity.

## Discussion

Our study underscores the significant influence of pH and LOI on shaping microbial community composition in soils from arable^28^ and permanent pasture fields, and demonstrates that the primary structuring factor for these microbial communities is pH. The association between soil pH and LOI at the national scale and across all habitat types showed a clear relationship between acidity and increasing LOI^29^. By focusing on highly managed soil systems situated on more basic parent material, we have constrained our study to soils that are under the most intensive forms of land use, where the conventional pH – LOI relationship is not consistently observed^26^. This study examined the underlying microbial mechanisms and niche dynamics in these soils, which may differ significantly from soils under less intensive land use or management strategies, such as woodlands, heaths, and bogs.

Our taxonomic analysis of the soils revealed a broad diversity, spanning several phyla, including abundant soil bacteria^30,31^ such as Actinobacteria, Proteobacteria, Planctomycetota, Chloroflexota, and Verrucomicrobiota. A positive correlation between pH and LOI with Entotheonellaeota, Crenarchaeota, and Acidobacteria suggests that these taxa may play critical roles in adapting to and mediating the effects of these environmental factors, providing further insight into the ecological strategies employed by these microbial communities. For example, certain taxa may be particularly well adapted to low-pH environments with high LOI, potentially through specialised metabolic pathways or ecological interactions that allow them to thrive under such conditions. Malik *et al*.^32^ previously reported taxa associated with low and high pH gradients with specific functional adaptations such as increased respiration and metabolism of aromatic compounds. Alternatively, microbial interactions, both mutual and antagonistic, may shape the composition and responses of the microbial community^33^. The role of soil carbon fluxes as a key function of microbial interactions^34^ is further supported by Garcia et al.^35^ who reported that soil carbon levels are influenced by microbial interactions. Furthermore, Cole et al.^36^ recently reported that in low-pH soils, land use intensification increases microbial physiological constraints and decreases carbon use efficiency. These findings contribute to a growing body of knowledge on the complex interactions between soil properties and microbial community structure, particularly in managed ecosystems.

The observed variability in microbial community composition and abundance across different soil types further led us to hypothesise that microbial genomic traits may reflect adaptive strategies. The positive correlation between genome size and rRNA copy number across these soils suggests that these genomic traits may be key indicators of microbial life history strategies, ecological adaptability, and evolutionary processes. Interestingly, we found that larger genomes and increased rRNA copies were associated with lower pH (see Table 1), suggesting soil-specific adaptations. Our observations are in agreement with the recent report by Wang et al.^9^, where genome sizes across soils from pH 3.7 to 7.2 decreased with higher pH, and also align with the findings of Malik et al.31, where at higher pH, a higher GC content is observed, leading to reduced genome size^37^. Our results suggest that non-acidophilic bacteria, typically observed in soil, do not follow the genome streamlining trends reported by Cortez et al.^38^. Larger genomes and higher rRNA copy numbers may provide a competitive advantage in nutrient-poor, complex, lower pH environments by enabling more efficient resource utilisation and rapid responses to environmental changes and opportunities. We hypothesise that this is likely because of lower pH soils being more environmentally and spatially variable. Although specific gene types are not associated with bacterial pH preferences^39^, anion/cation transporters, phosphatases, and efflux pumps have been identified in taxa that share pH associations^39^.

These results emphasise the importance of considering genomic traits when assessing microbial community dynamics in response to soil properties, as they offer valuable insights into the adaptive strategies of microbial taxa in different ecological contexts. Furthermore, mean sample SNB estimates revealed that pH was a major determinant of niche differentiation within the microbial community. It is likely that taxa in low [pH 5.5] to neutral pH soils are diverse^40^ and may possess broader ecological niches, potentially due to the need to exploit a wider range of resources or tolerate more variable conditions. This finding supports the idea that metabolic versatility, which is closely correlated with genome size, is higher in communities inhabiting low pH soils. As previously stated, Wilhelm et al.^10^ demonstrated a positive relationship between genome size and rRNA copy number in soil bacteria, however, unlike Wilhelm *et al*.^10^ our findings indicate that pH, rather than carbon content, is the dominant factor influencing genome size in the soils we examined. This suggests that care must be taken when using the average genome size as a proxy for carbon content or soil health, as pH may play a more significant role in shaping microbial genomic traits than previously thought. Additionally, Malik et al.^41^ identified a pH threshold of 6.2, above which significant shifts in carbon use efficiency occur, leading to smaller, more specialised genomes with narrower SNB at higher pH levels. The diversity in genome sizes coupled with SNB indicates that microbial taxa may employ different strategies to cope with the challenges posed by varying pH levels and significant additional drivers, such as genome size and rRNA copy number. Our results are in line with the hypothesis that larger genomes are associated with increased metabolic capacities^42^, potentially explaining the increased niche breadth. Collectively, our findings emphasise the need to understand how soil properties influence microbial ecology, particularly in managed ecosystems, where soil conditions may deviate from broader, natural trends.

Overall, this study provides new insights into the relationships among soil physicochemical properties, microbial community composition, and genomic traits in arable and permanent pasture soils. These findings challenge existing paradigms and highlight the need for further research on the specific mechanisms by which microbial communities adapt to and thrive in soils with atypical pH and LOI relationships. By integrating taxonomic, genomic, and ecological analyses, this study offers a comprehensive understanding of the factors that drive microbial diversity and function in these important agricultural ecosystems.

## Data and code availability

The code used for SNB analysis along with the Rscript used for figure generation with the associated files were uploaded to Zenodo. The files are publicly available and can be accessed via https://zenodo.org/records/13912858. Raw sequence files are available via the Sequence Read Archive. under BioProject ID PRJNA1168686.

## Contributions

TG: Methodology, formal analysis, validation, investigation, data curation, writing – original draft, visualisation, and project administration; SBB: methodology, formal analysis, validation, investigation, data curation, writing – original draft, and visualisation. RP: conceptualisation, resources, validation, project administration, funding acquisition, writing – review and supervision; BJ: resources, validation, and writing – review; RG: conceptualisation, resources, validation, project administration, funding acquisition, writing – review and supervision; AR: conceptualisation, project administration, and writing – review; MN: resources, validation, and writing – review; DSR: conceptualisation, resources, validation, project administration, funding acquisition, writing – original draft and supervision

## Acknowledgements and funding

This project was funded by UKCEH under the ASSIST programme (NERC Reference: NE/N018125/1). TG and DSR were also funded by the Natural Environmental Research Council (NERC) of UKRI via the PACIFIC (NE/X015947/1) and ‘Pushing the Frontiers’ MICRO- CYCLE (NE/Z000173/1) projects. SBB is supported by the Biotechnology and Biological Sciences Research Council (BBSRC), part of the UK Research and Innovation (UKRI), the Earlham Institute Strategic Programme Grant Decoding Biodiversity BBX011089/1, and its constituent work packages, BBS/E/ER/230002C (Decode WP3 Linking Fine-Scale Microbial Diversity to Ecosystem Functions).

The authors also thank the land management personnel, agronomists, and farmers involved in the project.

---

**Supplementary figure 1.**
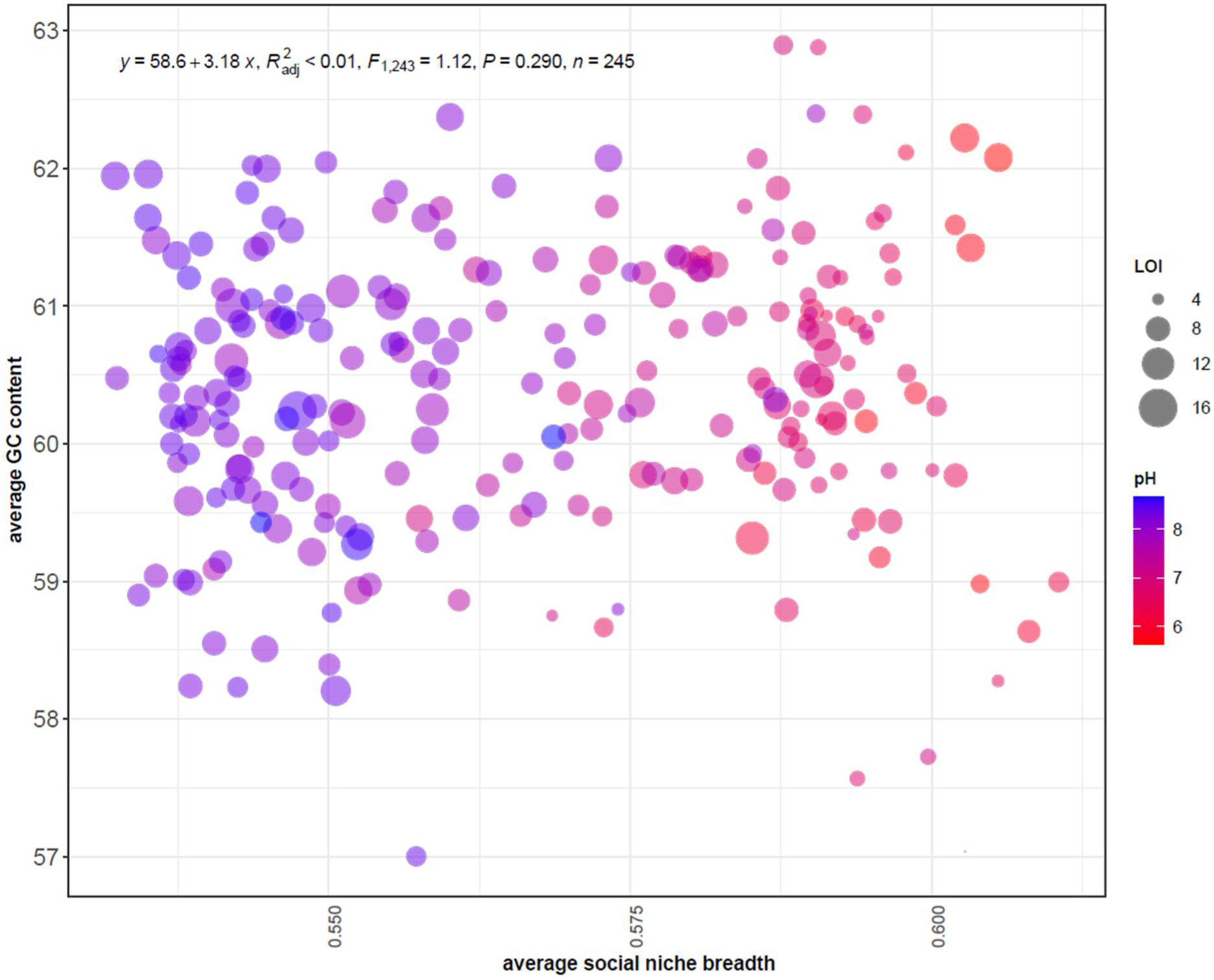
Scatter plot showing no relationship between weighted mean genome GC content and weighted mean social niche breadth, R^2^ = <0.01, *p* < 0.290, linear regression.

**Supplementary figure 2.**
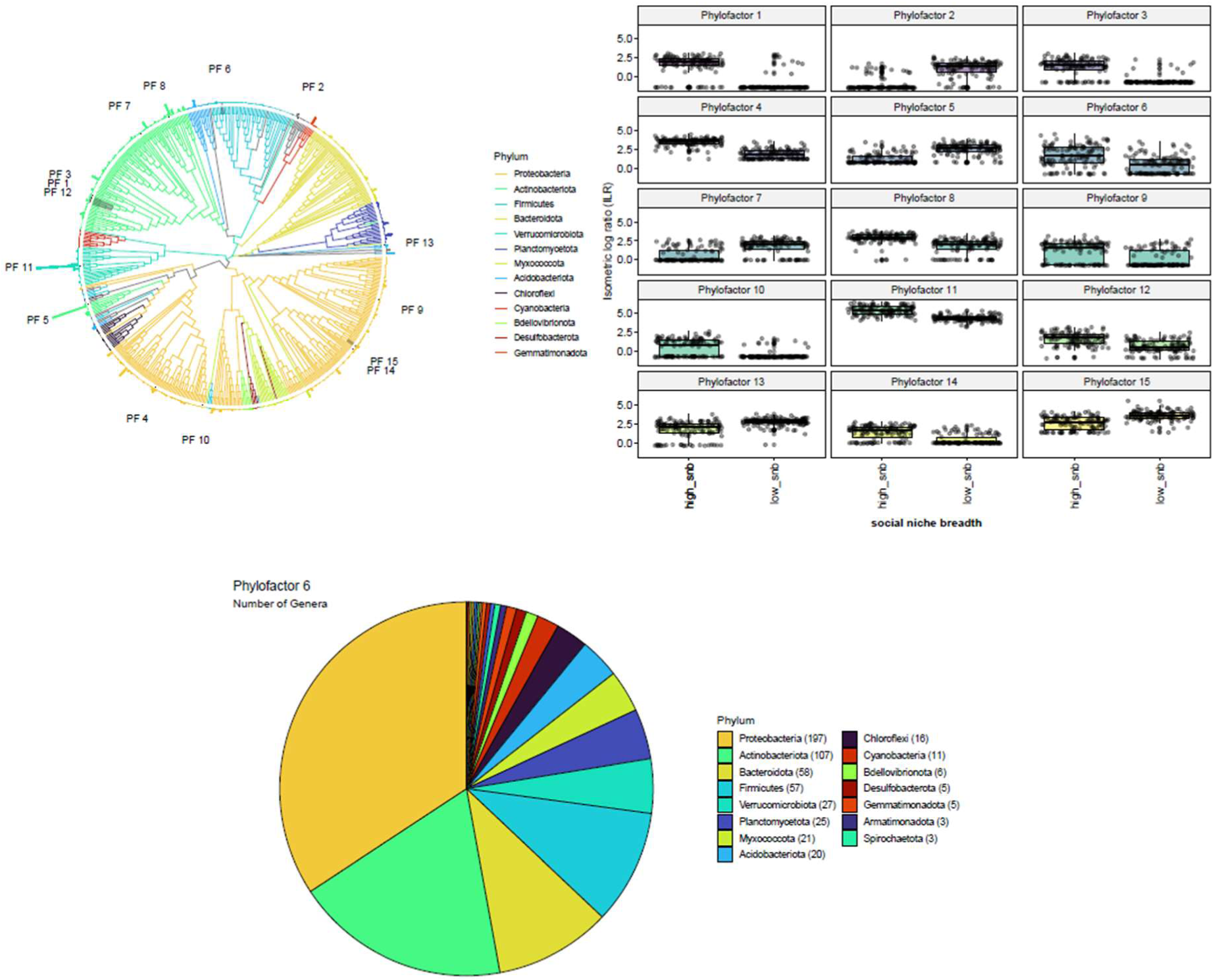
**(a)** PhyloFactor tree highlighting the factorial split (n=15) of the ASV-derived phylogenetic tree. Outer bars indicate the mean relative abundance of the individual ASVs across all samples. **(b)** Boxplots indicate the Isometric Log Ratio (ILR) derived from the phylofactor analysis, between ASVs with high-SNB and low-SNB scores and the 15 factors identified by PhyloFactor. **(c)** Representative phyla classification of PhyloFactor 6 indicating the influence of SNB irrespective of phylogeny and taxonomic bias.

**Supplementary figure 3.**
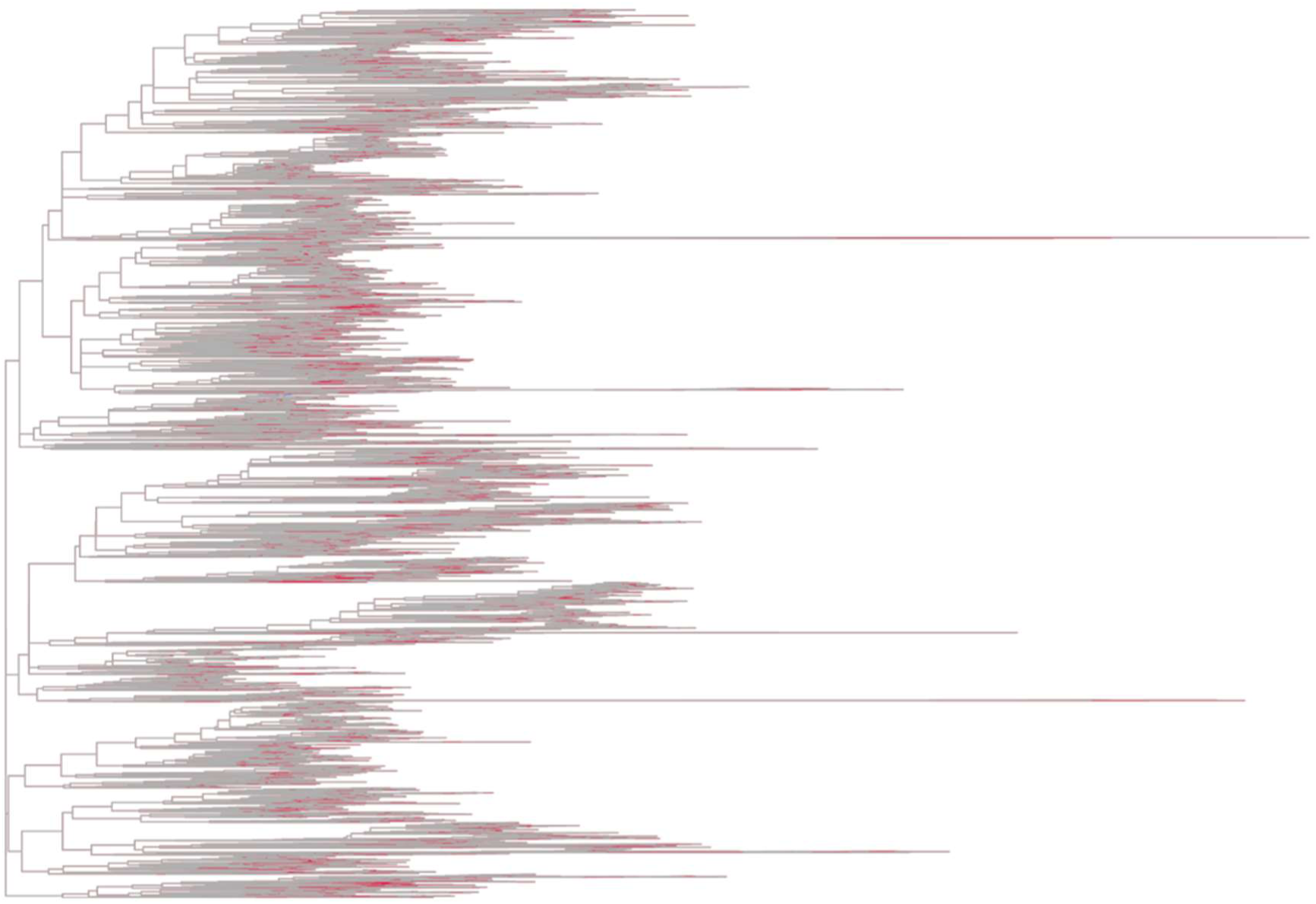
Phylogenetic tree highlighting the individual tips, i.e. ASVs, influenced by SNB, indicating a phylogenetic-independent signal for observed SNB scores.

## References

1 von Meijenfeldt, F. A. B., Hogeweg, P. & Dutilh, B. E. A social niche breadth score reveals niche range strategies of generalists and specialists. Nature Ecology & Evolution 7, 768–781 (2023). 10.1038/s41559-023-02027-7

2 Chuckran, P. F., Hungate, B. A., Schwartz, E. & Dijkstra, P. Variation in genomic traits of microbial communities among ecosystems. FEMS Microbes 2, xtab020 (2021). 10.1093/femsmc/xtab020

3. 3 DEFRA. Agricultural Land Use in United Kingdom at 1 June 2023, <https://www.gov.uk/government/statistics/agricultural-land-use-in-the-united-kingdom/agricultural-land-use-in-united-kingdom-at-1-june-2023> (2023).

4 Naylor, D., McClure, R. & Jansson, J. Trends in Microbial Community Composition and Function by Soil Depth. Microorganisms 10 (2022). 10.3390/microorganisms10030540

5 Zhang, K. et al. Interactive effects of soil pH and substrate quality on microbial utilization. European Journal of Soil Biology 96, 103151 (2020). 10.1016/j.ejsobi.2020.103151

6 Peng, Z. et al. Land conversion to agriculture induces taxonomic homogenization of soil microbial communities globally. Nature Communications 15, 3624 (2024). 10.1038/s41467-024-47348-8

7 Seaton, F. M., Griffiths, R. I., Goodall, T., Lebron, I. & Norton, L. R. Pasture age impacts soil fungal composition while bacteria respond to soil chemistry. *Agriculture*, Ecosystems & Environment 330, 107900 (2022). 10.1016/j.agee.2022.107900

8 Klappenbach, J. A., Dunbar, J. M. & Schmidt, T. M. rRNA operon copy number reflects ecological strategies of bacteria. Appl Environ Microbiol 66, 1328–1333 (2000). 10.1128/aem.66.4.1328-1333.2000

9 Wang, C. et al. Bacterial genome size and gene functional diversity negatively correlate with taxonomic diversity along a pH gradient. Nature Communications 14, 7437 (2023). 10.1038/s41467-023-43297-w

10 Wilhelm, R. C., Amsili, J. P., Kurtz, K. S. M., van Es, H. M. & Buckley, D. H. Ecological insights into soil health according to the genomic traits and environment-wide associations of bacteria in agricultural soils. ISME Commun 3, 1 (2023). 10.1038/s43705-022-00209-1

11 Carscadden, K. A. et al. Niche Breadth: Causes and Consequences for Ecology, Evolution, and Conservation. The Quarterly Review of Biology 95, 179–214 (2020). 10.1086/710388

12 Bennett, A. F. & Lenski, R. E. Evolutionary Adaptation to Temperature II. Thermal Niches of Experimental Lines of Escherichia coli. Evolution 47, 1–12 (1993). 10.2307/2410113

13 Kuang, J.-L. et al. Contemporary environmental variation determines microbial diversity patterns in acid mine drainage. The ISME Journal 7, 1038–1050 (2012). 10.1038/ismej.2012.139

14 Cobo-Simón, M. & Tamames, J. Relating genomic characteristics to environmental preferences and ubiquity in different microbial taxa. BMC Genomics 18, 499 (2017). 10.1186/s12864-017-3888-y

15 Walters, W. et al. Improved Bacterial 16S rRNA Gene (V4 and V4-5) and Fungal Internal Transcribed Spacer Marker Gene Primers for Microbial Community Surveys. mSystems 1, 10.1128/msystems.00009-00015 (2015). 10.1128/msystems.00009-15

16 Kozich, J. J., Westcott, S. L., Baxter, N. T., Highlander, S. K. & Schloss, P. D. Development of a dual-index sequencing strategy and curation pipeline for analyzing amplicon sequence data on the MiSeq Illumina sequencing platform. Appl Environ Microbiol 79, 5112–5120 (2013). 10.1128/aem.01043-13

17 Callahan, B. J. et al. DADA2: High-resolution sample inference from Illumina amplicon data. Nature Methods 13, 581–583 (2016). 10.1038/nmeth.3869

18 Martin, M. Cutadapt removes adapter sequences from high-throughput sequencing reads. 2011 17, 3 (2011). 10.14806/ej.17.1.200

19 Quast, C. et al. The SILVA ribosomal RNA gene database project: improved data processing and web-based tools. Nucleic Acids Res 41, D590–596 (2013). 10.1093/nar/gks1219

20 Liu, C., Cui, Y., Li, X. & Yao, M. microeco: an R package for data mining in microbial community ecology. FEMS Microbiology Ecology 97 (2020). 10.1093/femsec/fiaa255

21 Oksanen, J. et al. Vegan: Community Ecology Package. R Package Version 2.2-1 2, 1–2 (2015).

22 Markowitz, V. M. et al. IMG ER: a system for microbial genome annotation expert review and curation. Bioinformatics 25, 2271–2278 (2009). 10.1093/bioinformatics/btp393

23 Miao, J., Chen, T., Misir, M. & Lin, Y. Deep Learning for Predicting 16S rRNA Gene Copy Number. bioRxiv, 2022.2011.2026.518038 (2023). 10.1101/2022.11.26.518038

24 Washburne, A. D. et al. Phylogenetic factorization of compositional data yields lineage- level associations in microbiome datasets. PeerJ 5, e2969 (2017). 10.7717/peerj.2969

25 Kupka, D. & Gruba, P. Effect of pH on the sorption of dissolved organic carbon derived from six tree species in forest soils. Ecological Indicators 140, 108975 (2022). 10.1016/j.ecolind.2022.108975

26 Armbruster, M. et al. Bacterial and archaeal taxa are reliable indicators of soil restoration across distributed calcareous grasslands. European Journal of Soil Science 72, 2430–2444 (2021). 10.1111/ejss.12977

27 Griffiths, R. I. et al. The bacterial biogeography of British soils. Environmental Microbiology 13, 1642–1654 (2011). 10.1111/j.1462-2920.2011.02480.x

28 Aciego Pietri, J. C. & Brookes, P. C. Relationships between soil pH and microbial properties in a UK arable soil. Soil Biology and Biochemistry 40, 1856–1861 (2008). 10.1016/j.soilbio.2008.03.020

29. 29 Emmett, B. A. et al. Countryside Survey: Soils Report from 2007. Report No. 9/07, 192 (UK Centre for Ecology and Hydrology, 2010).

30 Bickel, S. & Or, D. The chosen few—variations in common and rare soil bacteria across biomes. The ISME Journal 15, 3315–3325 (2021). 10.1038/s41396-021-00981-3

31 Delgado-Baquerizo, M. et al. A global atlas of the dominant bacteria found in soil. Science 359, 320–325 (2018). doi:10.1126/science.aap9516

32 Malik, A. A., Thomson, B. C., Whiteley, A. S., Bailey, M. & Griffiths, R. I. Bacterial Physiological Adaptations to Contrasting Edaphic Conditions Identified Using Landscape Scale Metagenomics. mBio 8 (2017). 10.1128/mBio.00799-17

33 Ratzke, C. & Gore, J. Modifying and reacting to the environmental pH can drive bacterial interactions. PLoS Biol 16, e2004248 (2018). 10.1371/journal.pbio.2004248

34 Wu, H. et al. Unveiling the crucial role of soil microorganisms in carbon cycling: A review. Science of The Total Environment 909, 168627 (2024). 10.1016/j.scitotenv.2023.168627

35 García, F. C. et al. The temperature dependence of microbial community respiration is amplified by changes in species interactions. Nature Microbiology 8, 272–283 (2023). 10.1038/s41564-022-01283-w

36 Cole, L. et al. Land use effects on soil microbiome composition and traits with consequences for its ecosystem carbon use efficiency. bioRxiv, 2024.2004.2005.588235 (2024). 10.1101/2024.04.05.588235

37 Almpanis, A., Swain, M., Gatherer, D. & McEwan, N. Correlation between bacterial G+C content, genome size and the G+C content of associated plasmids and bacteriophages. Microb Genom 4 (2018). 10.1099/mgen.0.000168

38 Cortez, D., Neira, G., González, C., Vergara, E. & Holmes, D. S. A Large-Scale Genome-Based Survey of Acidophilic Bacteria Suggests That Genome Streamlining Is an Adaption for Life at Low pH. Front Microbiol 13, 803241 (2022). 10.3389/fmicb.2022.803241

39 Ramoneda, J. et al. Building a genome-based understanding of bacterial pH preferences. Science Advances 9, eadf8998 (2023). doi:10.1126/sciadv.adf8998

40 Luan, L. et al. Integrating pH into the metabolic theory of ecology to predict bacterial diversity in soil. Proceedings of the National Academy of Sciences 120, e2207832120 (2023). doi:10.1073/pnas.2207832120

41 Malik, A. A. et al. Land use driven change in soil pH affects microbial carbon cycling processes. Nature Communications 9, 3591 (2018). 10.1038/s41467-018-05980-1

42 Piton, G. et al. Life history strategies of soil bacterial communities across global terrestrial biomes. Nature Microbiology 8, 2093–2102 (2023). 10.1038/s41564-023-01465-0

